# High-resolution imaging atlas reveals context-dependent role of pancreatic sympathetic innervation in diabetic mice

**DOI:** 10.1101/2024.11.14.623708

**Authors:** Qingqing Xu, Yuxin Chen, Xinyan Ni, Hanying Zhuang, Shenxi Cao, Liwei Zhao, Leying Wang, Jianhui Chen, Wen Z Yang, Wenwen Zeng, Xi Li, Hongbin Sun, Wei L Shen

## Abstract

Gaining a better understanding of how sympathetic nerves impact pancreatic function is helpful for understanding diabetes. However, there is still uncertainty and controversy surrounding the roles of sympathetic nerves within the pancreas. To address this, we utilize high-resolution imaging and advanced three-dimensional (3D) reconstruction techniques to study the patterns of sympathetic innervation and morphology in islets of adult WT and diabetic mice. Our data shows that more than ∼30% α/β-cells are innervated by sympathetic nerves in both WT and diabetic mice. Also, sympathetic innervated α/β-cells are reduced in DIO mice, whereas sympathetic innervated β-cells are increased in *db/db* mice. Besides, in situ chemical pancreatic sympathetic denervation (cPSD) improves glucose tolerance in WT and *db/db* mice, but decreases in DIO mice. In situ cPSD also enhances insulin sensitivity in diabetic mice without affecting WT mice. Overall, our findings advance our comprehension of diabetes by highlighting the distinctive impact of pancreatic sympathetic innervation on glucose regulation.

**Highlights:** Pancreatic sympathetic innervation plays an important role in glucose homeostasis. In our study, we systematically investigate sympathetic innervation in the islets of WT and diabetic mice.

- 45.9% α-cells and 31.8% β-cells are innervated by sympathetic nerves in 10-week-old WT mice.
- Sympathetic innervated α/β-cells are reduced in DIO mice, whereas sympathetic innervated β-cells are increased in *db/db* mice.
- Unlike the core-mantle structure observed in WT and DIO mice, α-cells and β-cells are intermixed in *db/db* mice.
- Pancreatic sympathetic denervation elicits varied responses across diabetic models, where it improves glucose tolerance and insulin sensitivity in *db/db* mice, yet it increases insulin sensitivity but reduces glucose tolerance in DIO mice.

**Graphical abstract:** Summary of morphological and physiological phenotypes in diabetic mice.
All data are normalized to that of WT-10 mice. The numbers of “+” refer to their relative numbers. ↑ or ↓ refers to increase or decrease, ns refers to “not significant”. “−” refers to the experiment has not been carried out.

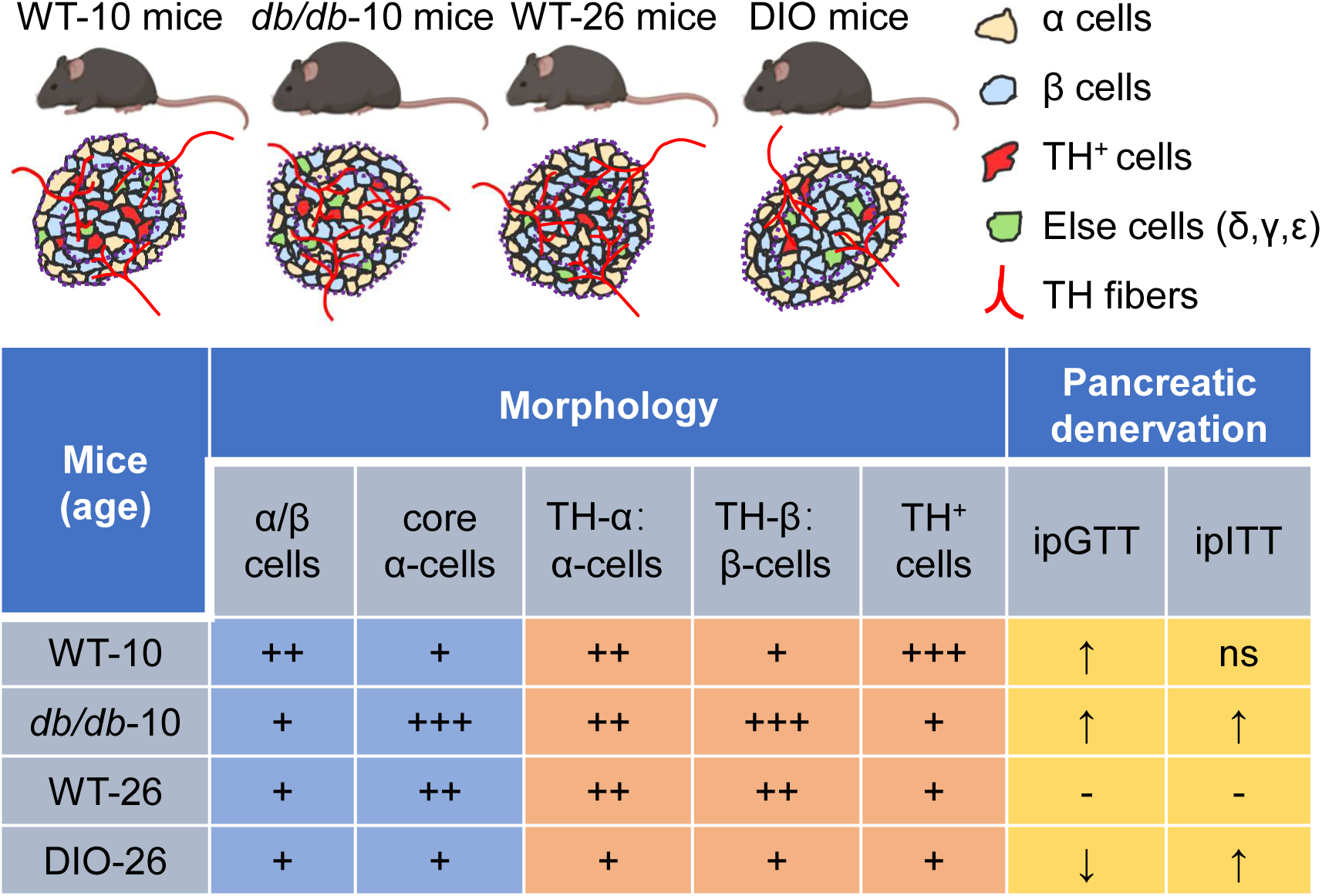

## Introduction

Glucose homeostasis is a critical physiological process that maintains the essential energy supply for the human body and prevents the detrimental effects of hyperglycemia or hypoglycemia [1]. This intricate regulation primarily hinges upon the secretion of hormones from the pancreatic islets [2]. Constituting a small fraction (1-2%) of the total pancreatic volume [3, 4], islets exhibit considerable variations in size [5]. The islets contain α-cells and β-cells that respectively secrete glucagon to increase blood glucose and insulin to lower them. These hormones work together to ensure that glucose levels remain within a narrow, healthy range, supporting the body’s energy needs. Changes in the histomorphology and function of islets correlate with the aberrant blood glucose levels [6]. Despite extensive research on the endocrine function of pancreas, our understanding of the intricate network of nerves that innervate pancreas and their roles in regulating blood glucose remains limited.

The autonomic nervous system, particularly the sympathetic nervous system, exerts its influence on the pancreas via visceral nerves originating from the prevertebral abdominal and superior mesenteric ganglia. This neural input is pivotal for the development and maturation of the pancreas [7–9]. Despite the acknowledged significance of sympathetic innervation in pancreatic function, the specific roles of these nerves within the pancreas remain unclear and controversial [10]. It is currently widely acknowledged that sympathetic innervation plays a predominant role in influencing the secretion of islet hormones through the neurotransmitter norepinephrine (NE). NE, released from postganglionic sympathetic fibers or the adrenal medulla, stimulates glucagon secretion by binding to β2-adrenergic receptors on α-cells and inhibits insulin secretion by binding to α2-adrenergic receptors on β-cells [11, 12]. Activation of the sympathetic nerves through electrical stimulation leads to the release of NE, which mimics the inhibitory effect on glucose-stimulated insulin secretion (GSIS) [13]. Similarly, administration of exogenous NE or adrenergic receptor agonists replicates the inhibitory effects of sympathetic nerve activation on GSIS [14]. Conversely, adrenergic receptor antagonists counteract the inhibitory effect of sympathetic nerve activation on insulin secretion [15]. In addition, activation of sympathetic nerves stimulates glucagon secretion and inhibit insulin release, leading to elevated blood glucose levels, which is essential for the ‘fight or flight’ response [16, 17]. However, it has been reported that the developmental loss of sympathetic nerves results in reduced insulin secretion and impaired glucose tolerance in adult mice [7]. To address the existing controversies, we need to manipulate the sympathetic nerves in pancreas to clarify their roles in the glucose metabolism. But, the detailed sympathetic innervation patterns within the pancreas also remain unclear. Previous study described that are few sympathetic fiber contacts on endocrine cells of islets in humans [18]. However, more recent applications of 3D imaging studies suggested significant sympathetic innervation in islets, particularly contacts on α- and δ-cells [19–21]. A view of the mouse pancreas, in which a rich supply of the sympathetic nerves was found to contact α-cells, but appear not to branch in space to establish contacts with β-cells [21, 22]. Impressively, Giannulis et al [23] found that islets of genetically obese mice showed increased sympathetic innervation of pancreatic islets, mainly with increased TH-positive fibers contacting β-cells. Nevertheless, multiple studies have suggested that exhibited a loss of sympathetic nerves in diabetes [24–27]. In light of these discordant findings, there is a pressing need to comprehensively map the pancreatic sympathetic innervation and elucidate its functional significance in glucose homeostasis.

In the present study, we systematically investigated the sympathetic innervation in pancreas of both wild type (WT) and diabetic mice. We observed that 45.9% α-cells and 31.8% β-cells are innervated by sympathetic nerves in 10-week-old WT mice. Moreover, sympathetic innervated α/β-cells were reduced in diet-induced obese (DIO) mice, whereas the sympathetic innervated β-cells were elevated in *db/db* mice. Interestingly, unlike the core-mantle structure observed in WT and DIO mice, α-cells and β-cells are intermixed in *db/db* mice. In addition, in situ cPSD led to an improvement in glucose tolerance in WT and *db/db* mice, but a decrement in DIO mice. Importantly, sympathetic denervation enhanced insulin sensitivity specifically in diabetic mice, with no observable effect on WT mice. These findings underscore the pivotal role of pancreatic sympathetic innervation in glucose homeostasis, providing new insights for deeply understanding diabetes.

## Materials and Methods

### Animals

Eight-week-old C57BL/6J mice and leptin-deficient *db/db* male mice were purchased from GemPharmatech Co., Ltd (Jiangsu, China). The DIO mice were established by feeding a high-fat diet (Rodent Diet With 60 kcal% Fat, D12492) to 10-week-old WT mice over a 16-week period. Mice were acclimated to standard temperature, humidity, and light conditions for 7 days before experiments. Mice were randomly assigned into 2 groups, containing vehicle group and 6-OHDA group. All animal experiments were conducted in strict adherence to the institutional protocols and ethical guidelines.

### Sympathetic nerve ablation

Mice (WT and *db/db* mice at 10-week-old, DIO mice at 26-week-old) were anesthetized with isoflurane. A feedback heater was used to keep mice warm during surgeries. The hair in the position of abdomen close to the pancreas was shaved, and the area was sterilized with 95% ethanol-soaked sterile gauze. A midline incision was made in targeted area of the abdomen skin (as viewed by the operator) to expose pancreas, and styptic powder was applied to the area to prevent bleeding. The pancreas was pulled out gently and fully exposed by sterile tweezers. The 6-OHDA (Sigma, H4381) was dissolved in saline solution containing 0.2% L-ascorbic acid (Sigma, A92902) to achieve a final concentration of 10 μg/μL. To pharmacologically ablate sympathetic nerves in pancreas, a 20-μL dose of 6-OHDA was evenly administered from the pancreatic head to the tail in the 6-OHDA group, utilizing a LEGATO^®^ 130 SYRINGE PUMP (#788130, RWD). An equivalent volume of 0.2% L-ascorbic acid solution was administered in the vehicle group. Then, the pancreas was gently placed back into original position. Mice were recovered in a warm blanket before they were transferred to housing cages. One to two weeks post-pharmacological ablation, the mice were employed for subsequent experimental procedures.

### Intraperitoneal glucose tolerance tests (GTTs) and intraperitoneal insulin tolerance tests (ITTs)

GTTs were conducted in mice after 6-h fasting following an intraperitoneal (ip) injection of glucose (20% dextrose) at a dose of 2.0 g/kg body weight. Blood samples were collected from the mouse orbital vein at the indicated time for subsequent analysis via the Enzyme-Linked Immunosorbent Assay (ELISA). ITTs were performed in 6-h fasted mice by injecting either 0.75 units/kg body weight (for WT mice) or 1.0 units/kg (for DIO mice) or 1.5 units/kg (for *db/db* mice) recombinant Human Insulin Injection (Novo Nordisk, 100U/mL, China).

### Physiological measurements

Glucose measurement was determined by a hand-held glucometer (Accu-Chek Performa Connect, Roche, Switzerland). Blood samples were collected from the tail vein in ad-lib fed mice. Concentrations of serum insulin were measured by Insulin Elisa kit (J&L Biology, #JL1145) according to the manufacturer’s instruction.

### Immunohistochemistry

Mice were anesthetized and perfused with Phosphate-Buffered Saline (PBS) followed by 4% Paraformaldehyde (PFA). Pancreas was excised, post-fixed overnight. The pancreas was dehydrated sequentially in 15% sucrose solution and 30% sucrose solution for 2-day at 4°C, then sectioned at 40-μm thicknesses on a cryostat microtome (Leica, CM3050s). The slices were washed three times in PBST (PBS with 0.3% Triton X-100, v/v), and blocked with QuickBlock™ blocking buffer for 2-h at room temperature. Then, slices were incubated 36-h with primary antibodies (1:500) diluted in QuickBlock™ primary antibody dilution buffer. After that, the slices were washed 3 times in PBST before being incubated in secondary antibodies (1:1000) for 12-h at room temperature. Pancreatic sections were washed 3 times in PBST and cover-slipped with DAPI for subsequent image processing. The immunohistochemical reagents utilized in this study are detailed in Supplementary Table 1.

### Imaging processing and analysis

High resolution imaging was performed using Leica SP8 STED 3X confocal microscopes (operated with Leica Application Suite X, version 3.5.2). Excitation was delivered using 405 nm, 488 nm, 568 nm and 633 nm laser lines. Signals were detected at 410-481 nm (DAPI), 498-560 nm (Alexa 488), 578-630 nm (Alexa 568) and 641-739 nm (Alexa 633) using HyD spectral detectors. Confocal images (pinhole = airy 1, step size = 0.5-μm) of randomly selected islets (9-15 islets per section) were acquired on a Leica SP8 confocal microscope. Quantitative analysis was calculated using Imaris (9.7.2) and Fiji open-source software. A subset of experiments was performed using a Nikon CSU-W1 Sora 2 Camera confocal equipped with a 20× 0.8 / air objective.

To calculate numbers of β-cells and α-cells per slice, each islet was evaluated to obtain the cell numbers within this islet positive for insulin and glucagon using Imaris versions 9.7.2 (Bitplane AG, Zürich, Switzerland). Imaris software was used to create digital surfaces covering the islets and innervation to automatically determine volumes and intensity data. Volume reconstructions were performed via the surface function with local contrast background subtraction. For detection of islets, the threshold factor was set to “20” corresponded to the largest α- and β-cell diameter in each sample, and used the automated surface algorithm with a 5-µm “smooth texture” for excluding hypointense (non-tissue filled) regions. Numbers of islets with core α-cells were determined according to a rule reported previously [7]: using the edge of the islet as the boundary, indent 2 layers of cells toward the interior of the islet, and quantify the number of α-cells within the circle.

The nerve fiber plexus was reconstructed in 3D stacks of images. For detection of nerves, the threshold factor was set to 1.5-μm. In confocal images, digital surfaces were created to cover nerve fibers and individual β-cell and α-cell. The Imaris Distance Transform Matlab XTension function was used to calculate the distance of each α- and β-cell surface from the innervation surface. This measurement was subsequently employed to determine the distance between sympathetic nerves and individual α- or β-cell with a distance of 0 indicating a nerve contact [20, 28].

### Morphometric measurements

For quantification, slices were digitized with OlyVIA Vs120 software on a fluorescence scan at 20X magnification. Quantitative analysis of islet area was performed with Image J software using the positive-pixel count algorithm, expressed as μm^2^. When analyzed the islet size, we referred to the classification methods [29]. The classification criteria for islets are specified as follows: Islets with an area exceeding 25,000 μm² are categorized as ‘Large Islets’ (LI), those with an area less than 10,000 μm² are categorized as ‘Small Islets’ (SI), and those in between are categorized as ‘Medium Islets’ (MI). To determine the sizes of islets and counts of α/β-cells, immunostaining of insulin and glucagon was performed in pancreatic sections. This analysis involved manually capturing images and counting the islets. Image analysis was employed to quantify the entire section, followed by the identification of insulin/glucagon-positive areas using Image J software to calculate the islet area. The ‘closed polygon’ tool in Image J software was utilized to determine the parameters of the islets.

### Quantification and statistical analysis

All statistical data from individual segmented islets were exported from Imaris as to Excel^®^ (Microsoft^®^, version 2010), and each α- or β-cell received an individual ID for sorting purposes. All results are represented as mean ± SEM for the indicated number of observations. Student’s t-test was used for two-group comparisons, one-way ANOVA was used for multi-groups comparisons and two-way ANOVA with multiple comparisons was used for groups mixed by time factorial designs. Data were analyzed using Prism 8.0. A p value <0.05 was considered as statistically significant.

## Results

### Distribution of sympathetic nerves in wild-type pancreatic islets

To characterize the distribution of sympathetic nerves in the pancreas, we performed the immunostaining for insulin (Ins), glucagon (Gcg) and tyrosine hydroxylase (TH), which indicate β-cells, α-cells and sympathetic nerves respectively (Figure. 1A). To further explore the detailed spatial distribution, we reconstructed these confocal images (Figure. 1B). Considering the different sizes, we categorized islets into three groups: small islets (SI, area ≤ 10,000 μm^2^), medium islets (MI, 10,000 - 25,000 μm^2^) and large islets (LI, ≥ 25,000 μm^2^) (Figure. 1A). We found that there were 58.7% SI, 17.5% MI and 23.8% LI in adult WT mice (10 weeks old, Figure. 1C). To better understand the distribution of α-cells in islets, we used the edge of islet as the boundary, indented 2 layers of cells toward the interior, and quantified the number of α-cells within the circle. Typically, unlike β-cells, which are ubiquitously expressed throughout all islets, most α-cells are within the shell of SI, MI, and LI (Figure. 1A and 1D). We also quantified the numbers of α/β-cells and found that the ratio of α-: β-cells did not change among different sizes of islets (Figure. 1E). Based on the 3D architecture, we classified α/β-cells into TH-innervated ones and un-innervated ones (Figure. 1B, Supplementary Movie S1-S4). We found that 60.9% α-cells are innervated by TH in SI, 45.3% in MI and 29.5% in LI (Figure. 1F).

**Figure 1.**
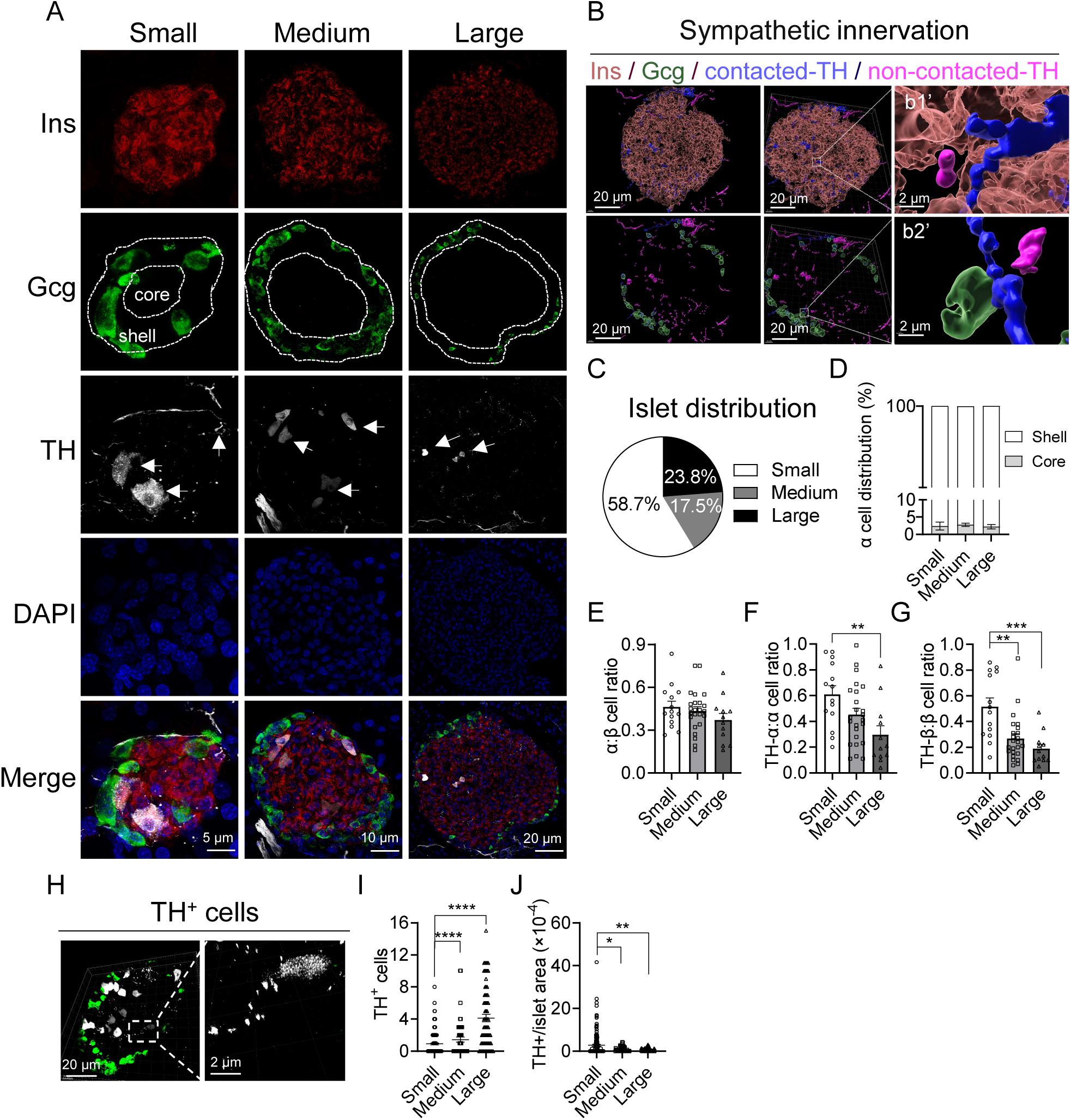
Distribution of sympathetic nerves in the wild-type pancreatic islets. (A) Immunostaining for Insulin (Ins, red), Glucagon (Gcg, green), and Tyrosine hydroxylase (TH, gray) in WT pancreas. The islets exhibit a uniform architecture with α-cells at the islet periphery surrounding a β-cell core. It is about two cell layers between the core and shell dashed line. White arrows indicate resident TH^+^ cells in islets. (B) Schematic diagram of 3D reconstruction. β-cells (pink, transparent), α-cells (green, transparent), α/β-cells contacted TH (blue, opaque), α/β-cells uncontacted TH (purple, opaque). (C) Distribution of islets in different sizes (Small islets, n=168; medium islets, n=51; large islets, n=70). (D) Distribution of α-cells located in the core and shell of islets, defined in (A). (E) The ratio of α to β cell numbers. (F-G) Percentage of TH-innervated α-cells out of total α-cells (F) and TH-innervated β-cells out of total β-cells (G). (H) Representative images of resident TH^+^ cells. (I-J) Distribution of TH^+^ cells in different islets (I) and normalized distribution using islet area (J). Data are mean ± SEM and analyzed by one-way ANOVA. *p< 0.05, **p< 0.01, ***p< 0.001, ****p < 0.0001.

Similarly, 51.6% β-cells were innervated by TH in SI, 26.7% in MI and 18.6% in LI (Figure. 1G). Interestingly, we found there were some TH positive (TH^+^) cells within islets (Figure. 1H). Meanwhile, the total number of these cells increased from SI to LI (Figure. 1I). Yet, when normalized to islet areas, the relative number of these cells decreased from SI to LI (Figure. 1J). Overall, we found that the pancreas of WT mice was predominantly composed of smaller islets and most α-cells were preferentially distributed within the shell of islets. Notably, we observed an inverse correlation between the size of islets and the proportion of α/β-cells innervated by sympathetic nerves, suggesting that the degree of sympathetic innervation per α/β-cell may be attenuated with islet enlargement, which could have functional implications for islet endocrine activity and glucose regulation.

### Pancreatic sympathetic nerves were reduced in DIO mice

To evaluate whether sympathetic innervation is altered in diabetic mice, we induced DIO models by feeding C57 mice a high fat diet (HFD) (Figure. 2A). Compared to WT-26 mice (WT mice at 26 weeks old), we found that islet sizes increased significantly in DIO-26 mice (DIO mice at 26 weeks old with 16 weeks HFD) (Figure. 2B, Supplementary Figure S1A and S1B). Compared to WT-10 mice (WT mice at 10 weeks old), WT-26 mice exhibited reduced SI percentage (WT-26: WT-10 = 52.9%: 58.7%) and increased MI percentage (WT-26: WT-10 = 24.0%: 17.5%) respectively (Figure. 2C). In comparison with WT-26 mice, DIO mice showed decreased SI percentage (DIO: WT-26 = 48.6%: 52.9%), MI percentage (DIO: WT-26 = 22.1%: 24.0%) and increased LI percentage (DIO: WT-26 = 29.3%: 23.1%) (Figure. 2C). Similar as observed in WT-26 mice, most α-cells were located within islet shells in DIO mice (Figure. 2D). Moreover, the ratio of α-: β-cells was significantly reduced with aging in WT mice, but no difference was found between WT-26 and DIO-26 (Figure. 2E). We also found no significant change in TH-innervated α-cells between WT-10 (45.9%) and WT-26 mice (48.5%), whereas it was significantly decreased in DIO mice (37.9%) compared to WT-26 mice (48.5%) (Figure. 2F). However, we found a significant increase in TH-innervated β-cells of WT-26 mice (46.1%) compared to WT-10 mice (31.8%) (Figure. 2G). Meanwhile, it was distinctly decreased in DIO mice (34.1%) compared to WT-26 mice (46.1%) (Figure. 2G). In addition, the normalized number of TH^+^ cells was reduced in WT-26 or DIO-26 mice compared to corresponding control groups (WT-26 vs WT-10, DIO-26 vs DIO-10), whereas there was no significant difference in DIO mice and WT-26 mice (Figure. 2H).

**Figure 2.**
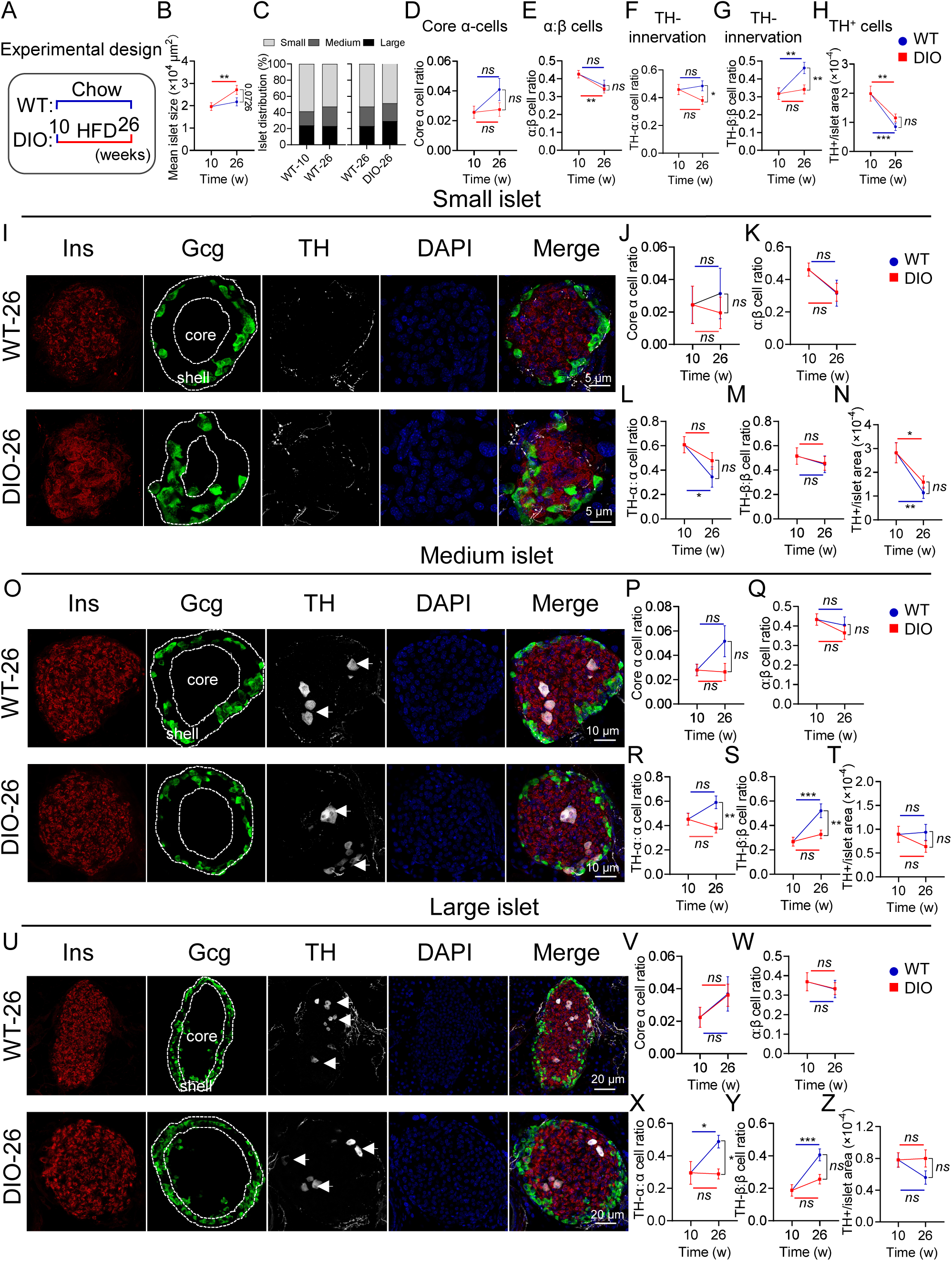
Distribution of sympathetic nerves in pancreatic islets of DIO mice. (A) Experimental design for DIO mice (WT-10/26 refers to WT mice of 10 weeks old and 26 weeks old respectively, DIO-26 refers to WT-10 mice fed with HFD for 16 weeks). (B-C) Comparison of mean islet area (B) and islet distribution (C) between WT and DIO mice (WT-10, n=289 from 4 mice; WT-26, n=267 from 4 mice; DIO-26, n=427 from 5 mice). (D) Percentage of α-cells located in the core of islets. All size of islets is quantified. The boundaries are defined in (I), (O) and (U). (E) The ratio of α to β cell numbers for all islets. (F-G) Percentage of TH-innervated α-cells out of total α-cells (F) and TH-innervated β-cells out of total β-cells (G). (H) Quantification of TH^+^ cells in all islets. (I) Representative images of small islets in WT-26 mice and DIO-26 mice (WT-10, n=14; WT-26, n=13; DIO-26, n=17). (J) Percentage of α-cells located in the core of small islets. (K) The ratio of α to β cell numbers in small islets. (L-M) Percentage of TH-innervated α-cells out of total α-cells (L) and TH-innervated β-cells out of total β-cells (M) in small islets. (N) Quantification of TH^+^ cells in small islets. (O-T) Representative images and quantifications for medium islets, with similar quantification as (I-N) (WT-10, n=24; WT-26, n=17; DIO-26, n=24). (U-Z) Representative images and quantifications for large islets, with similar quantification as (I-N) (WT-10, n=12; WT-26, n=15; DIO-26, n=19). Data are mean ± SEM and analyzed by unpaired t test. *p < 0.05, **p < 0.01, ***p< 0.001, ****p < 0.0001.

In SI (Figure. 2I), the majority of α-cells were located within islet shells in both WT and DIO mice (Figure. 2J). We also found that there was no difference in the ratio of α-: β-cells (Figure. 2K), TH-innervated α-cells (Figure. 2L), TH-innervated β-cells (Figure. 2M) and the normalized number of TH^+^ cells (Figure. 2N) between DIO and WT mice. In MI (Figure. 2O), the distribution of α-cells in WT and DIO mice was similar as that in SI (Figure. 2P). Meanwhile, there was also no difference in the ratio of α-: β-cells between DIO and WT mice (Figure. 2Q). Besides, the TH-innervated α-cells were obviously decreased in DIO mice (38.0%) compared to WT-26 mice (59.0%) (Figure. 2R). So were the TH-innervated β-cells (DIO: WT-26 = 32.7%: 52.0%) (Figure. 2S). However, there was no difference in the normalized number of TH^+^ cells (Figure. 2T). In LI (Figure. 2U), most α-cells were also located within islet shells in both WT and DIO mice (Figure. 2V). And there was no difference in the ratio of α-: β-cells between DIO and WT mice (Figure. 2W). Furthermore, the TH-innervated α-cells were decreased in DIO mice (28.8%) compared to WT-26 mice (48.8%) (Figure. 2X). So were the TH-innervated β-cells (DIO: WT-26 = 25.6%: 40.6%, Figure. 2Y). Similar to SI and MI, there was no difference in the normalized number of TH^+^ cells between DIO and WT mice (Figure. 2Z).

Taken together, we concluded that islets undergo progressive enlargement with aging and elevated blood glucose levels. Moreover, HFD decreased the sympathetic innervation on both α-cells and β-cells.

### Sympathetic-innervated α/β-cells altered in *db/db* mice

We further examined genetically diabetic *db/db* mice at 10 weeks old (db-10) and 26 weeks old (db-26). The mean islet size of *db/db* mice was remarkably larger than that of WT mice, especially in SI and LI (Figure. 3A, Supplementary Figure S2A and S2B). Compared to WT-10 mice (58.7% for SI, 17.5% for MI and 23.8% for LI), SI percentage (25.0%) in db-10 mice was drastically reduced, MI percentage (31.6%) and LI percentage (43.4%) were increased (Figure. 3B and S2C). In comparison with WT-26 mice (52.9% for SI, 24.0% for MI and 23.1% for LI), db-26 mice showed significant reduced in SI percentage (20.1%) and increased in LI percentage (59.9%), but no difference in MI percentage (20.0%) (Figure. 3B and S2C). Unlike islets with a core–mantle structure in WT mice (Figure. 1A), α-cells and β-cells are intermingled in *db/db* mice (Figure. 3C). In contrast to WT mice, α-cells in *db/db* mice exhibited a random distribution across the islet area, rather than being confined to the shell (Figure. 3C). Besides, the ratio of α:β cells was significantly decreased in db-10 mice compared to WT-10 mice, but no difference between db-26 and WT-26 mice (Figure. 3D). Intriguingly, there was no significant difference in TH-innervated α-cells between WT-10 (45.9%) and db-10 mice (45.3%), whereas it was obviously decreased with aging in *db/db* mice, and its proportion in db-26 mice (31.9%) is significantly lower than WT-26 mice (48.5%) (Figure. 3E). However, we found that TH-innervated β-cells were increased in db-10 mice (66.2%) compared to WT-10 mice (31.8%). But there was no difference between db-26 (52.0%) and WT-26 (46.1%) (Figure. 3F). Moreover, the normalized TH^+^ cells was decreased in *db/db* mice compared to WT mice regardless of age (Figure. 3G).

**Figure 3.**
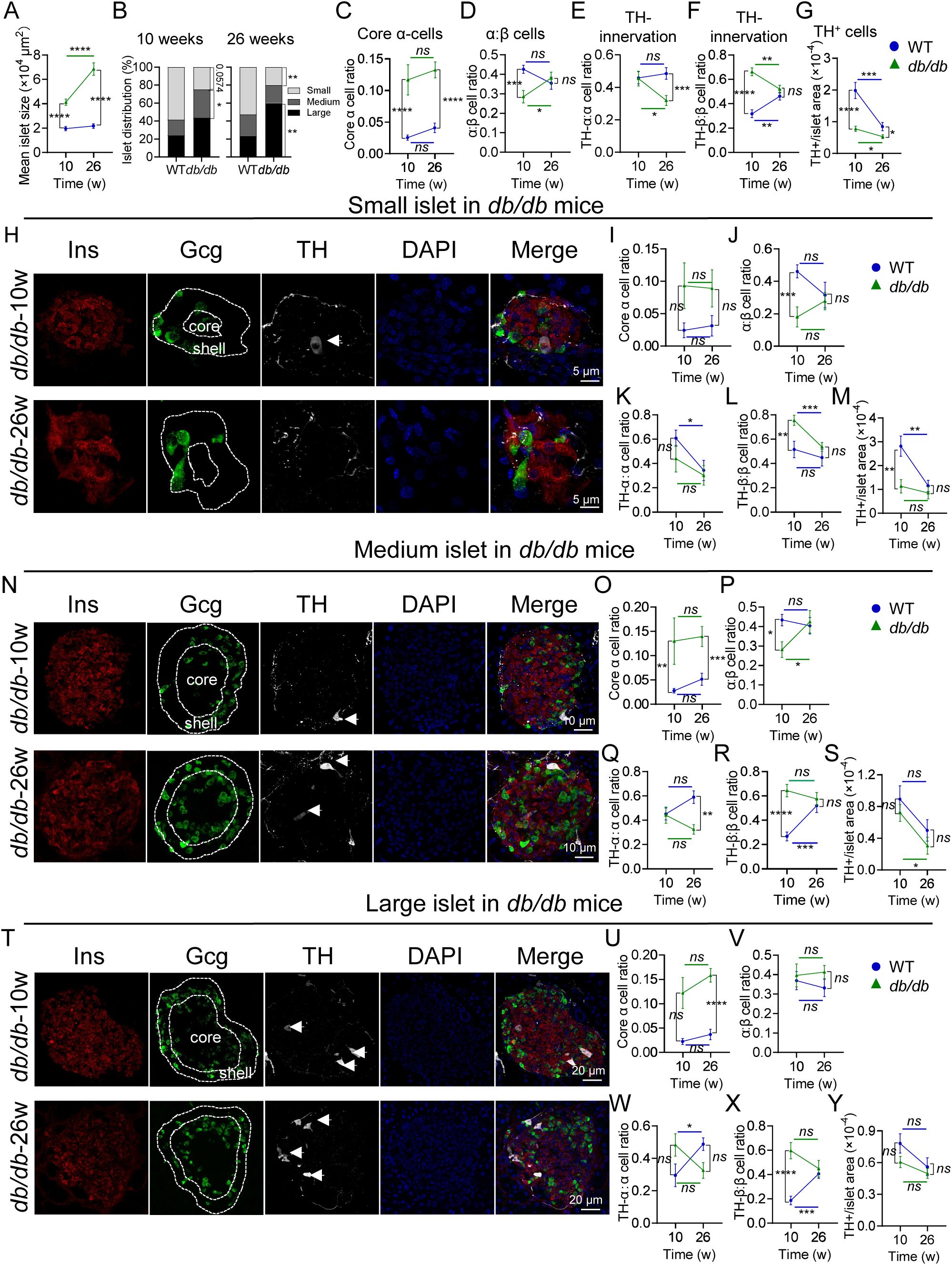
Distribution of sympathetic nerves in pancreatic islets of *db/db* mice. (A-B) Average area and distribution of islets with different sizes in WT mice and *db/db* mice (WT-10, n=289 from 4 mice, WT-26, n=267 from 4 mice; db-10, n=390 from 3 mice, db-10, n=256 from 3 mice). (C) Percentage of α-cells located in the core of islets. (D) The ratio of α to β cell numbers for all islets. (E-F) Percentage of TH-innervated α-cells out of total α-cells (E) and TH-innervated β-cells out of total β-cells (F), in all islets. (G) Quantification of TH^+^ cells. (H) Representative images of small islets in *db/db* mice (WT-10, n=14; WT-26, n=13; db-10, n=10; db-26, n=10). (I) Percentage of α-cells located in the core of small islets. (J) The ratio of α to β cell numbers for small islets. (K-L) Percentage of TH-innervated α-cells out of total α-cells (K) and TH-innervated β-cells out of total β-cells (L) in small islets. (M) Quantification of TH^+^ cells. (N-S) for medium islets, with similar quantification as in (H-M) (WT-10, n=24; WT-26, n=17; db-10, n=15; db-26, n=14). (T-Y) for large islets, with similar quantification as in (H-M) (WT-10, n=12; WT-26, n=15; db-10, n=10; db-26, n=13). Data are mean ± SEM and analyzed by unpaired t test. *p < 0.05, **p < 0.01, ***p< 0.001, ****p < 0.0001.

In SI (Figure. 3H), most α-cells distributed within the core of islets in *db/db* mice (Figure. 3I). The ratio of α-: β-cells was significantly decreased in db-10 mice compared to WT-10 mice, and no difference between db-26 and WT-26 mice (Figure. 3J). TH-innervated α-cells were decreased with aging in both WT (WT-26: WT-10 = 34.4%: 60.9%) and *db/db* mice (db-26: db-10 = 30.2%: 44.0%). But there was no difference between WT and *db/db* mice (Figure. 3K). TH-innervated β-cells also decreased with aging in both WT mice (WT-26: WT-10 = 44.8%: 51.6%) and *db/db* mice (db-26: db-10 = 53.7%: 75.5%) (Figure. 3L). Obviously, it was significantly higher in *db/db* mice (Figure. 3L). The normalized TH^+^ cells were decreased in db-10 mice compared to WT-10 mice (Figure. 3M). In MI (Figure. 3N), a notable increase in the distribution of α-cells was observed within the core of islets in *db/db* mice compared to WT mice (Figure. 3O). Moreover, the ratio of α-: β-cells was significantly decreased in db-10 mice compared to WT-10 mice, and no difference between db-26 and WT-26 mice (Figure. 3P). Although there was no difference between db-10 (44.3%) and WT-10 mice (45.3%), the TH-innervated α-cells were decreased in db-26 mice (32.4%) compared to WT-26 mice (59.0%) (Figure. 3Q). TH-innervated β-cells were increased in db-10 mice (64.4%) compared to WT-10 mice (26.7%), but no difference between db-26 mice (57.5%) and WT-26 mice (52.0%) (Figure. 3R). In addition, there was no difference in the normalized TH^+^ cells between *db/db* mice and WT mice (Figure. 3S). In LI (Figure. 3T), more α-cells distributed in the core of islets in db-26 mice compared to WT-26 mice (Figure. 3U). However, there was no difference in the ratio of α-: β-cells (Figure. 3V), TH-innervated α-cells (Figure. 3W) and the normalized TH^+^ cells (Figure. 3Y) between *db/db* and WT mice. Interestingly, TH-innervated β-cells were increased in db-10 mice (59.7%) compared to WT-10 mice (18.75%), but no difference between db-26 mice (44.8%) and WT-26 mice (40.6%) (Figure. 3X). In summary, we concluded that islets in *db/db* mice exhibited a random distribution of α-cells and β-cells compared to those in WT mice which have a uniform structure. Besides, there are more sympathetic innervated β-cells in *db/db* mice. All these data suggested that the distribution of α/β-cells and their sympathetic innervation may be involved in the development of diabetes.

### Effect of cPSD on the glucose metabolism

To examine the role of sympathetic innervation in glucose metabolism, we first executed chemical denervation in the pancreas of WT mice in situ. To achieve this, we administered 6-Hydroxydopamine (6-OHDA) into the pancreas, aiming to ablate the sympathetic nerves (Figure. 4A). The TH expression was significantly reduced after 25 days following 6-OHDA administration (Figure. 4B and 4C). Although there was no change in body weight (Figure. 4D), glucose tolerance was improved after 15 days following 6-OHDA injection, as determined by the intraperitoneal glucose tolerance test (GTT) (Figure. 4E). We also measured insulin (Figure. 4F) and glucagon (Figure. 4G) concentration in the GTT, and observed a significant elevation in the accumulation of insulin (Figure. 4F), implying sympathetic denervation increases insulin secretion. However, sympathetic denervation had no effect on the insulin sensitivity in WT mice as measured by intraperitoneal insulin tolerance test (ITT) (Figure. 4H).

**Figure 4.**
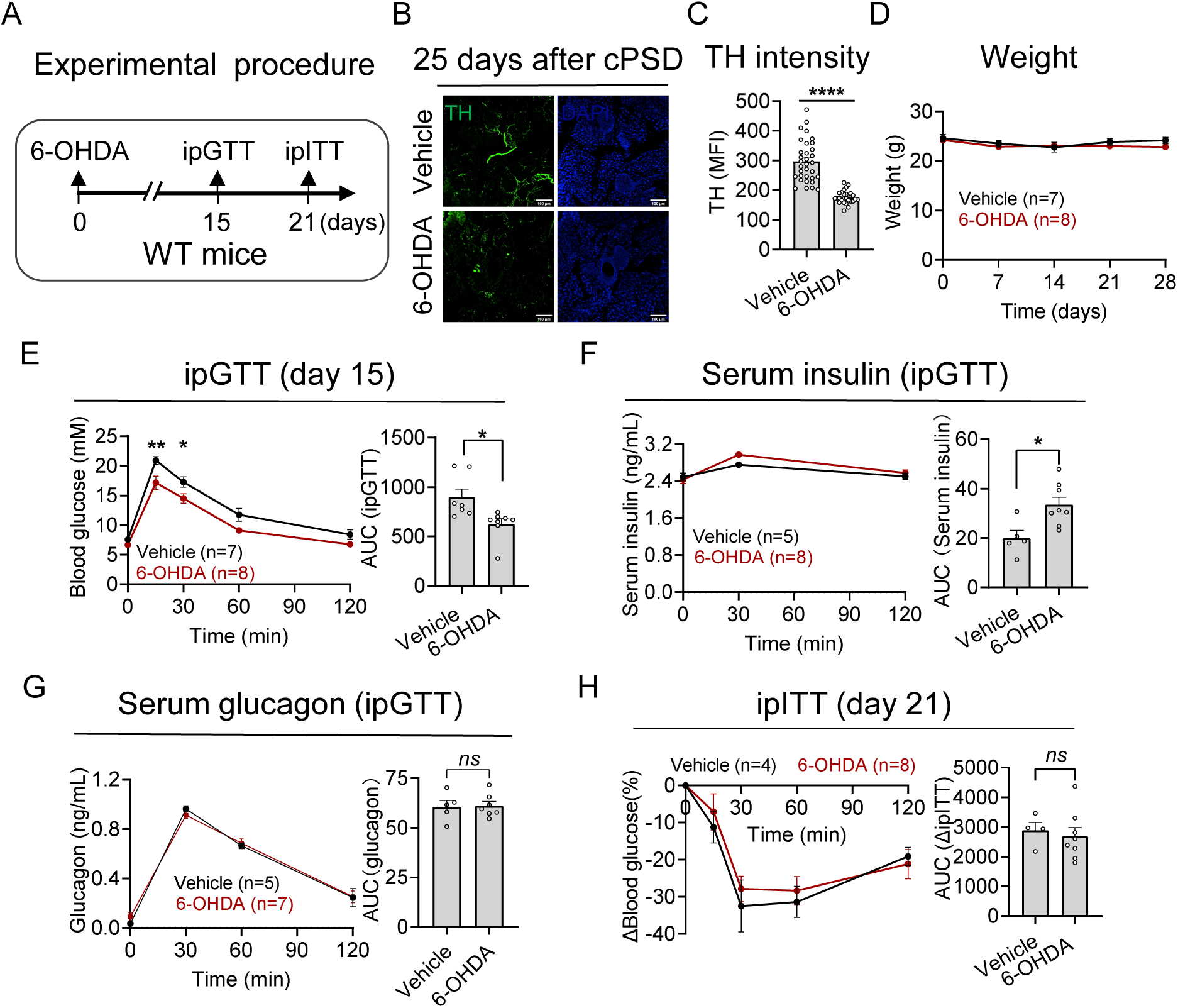
Effects of pancreatic sympathetic denervation in WT mice. (A) An experimental design was implemented to examine the impact of sympathetic denervation on WT mice (10-week-old), involving multiple injections of 6-OHDA administered at various sites within the pancreas. (B) Immunofluorescence of TH in pancreas after 25 days following 6-OHDA injection. (C) Quantification of the mean fluorescence intensity (MFI) (normalized to vehicle group) of TH. Scale bar, 100 μm. (D) Changes of body weight after sympathetic denervation. (E) Changes of blood glucose (left) and area under curve (AUC, right) of WT mice subjected to intraperitoneal glucose tolerance test (ipGTT). (F) Changes of serum insulin (left) and AUC (right) during ipGTT. (G) Changes of serum glucagon (left) and AUC (right) during ipGTT. (H) Changes in blood glucose (left) and AUC (right) of WT mice subjected to intraperitoneal insulin tolerance test (ipITT). All data are mean ± SEM and analyzed by two-way ANOVA with multiple comparisons and unpaired t test. *p < 0.05, **p < 0.01, ****p < 0.0001, ns, p > 0.05.

Next, we injected 6-OHDA into the pancreas of DIO mice (Figure. 5A). We found there was no difference in body weight or blood glucose between 6-OHDA and vehicle group (Figure. 5B and 5C). However, the glucose tolerance was deteriorated after 6-OHDA injection (Figure. 5D). Meanwhile, the accumulation of insulin secretion was reduced and glucagon was increased in the GTT compared to the vehicle group (Figure. 5E and 5F). Furthermore, DIO mice exhibited increased insulin sensitivity after 21 days following 6-OHDA injection as determined by ITT (Figure. 5G).

**Figure 5.**
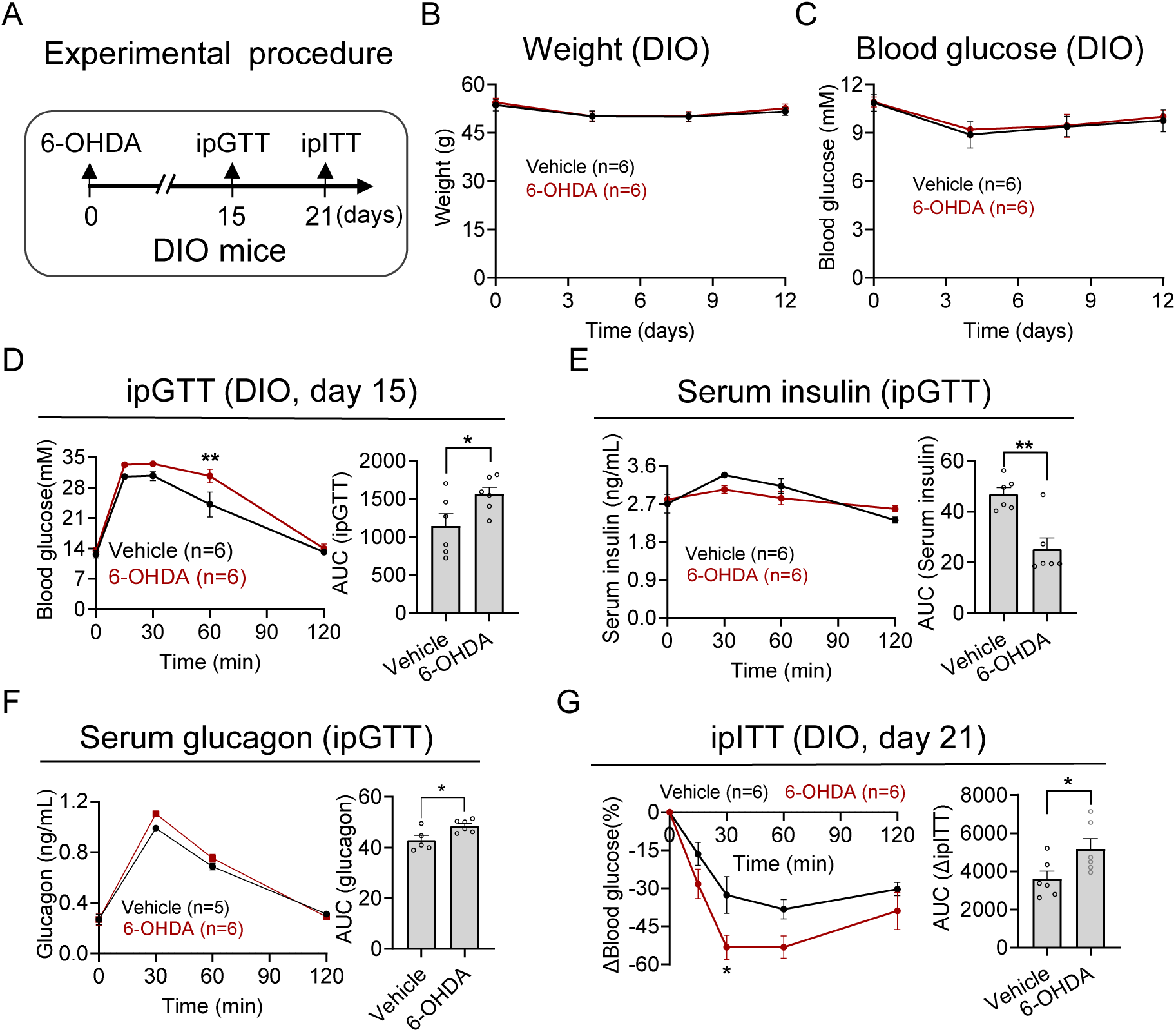
Effects of pancreatic sympathetic denervation in DIO mice. (A) An experimental design was implemented to examine the impact of sympathetic denervation on DIO mice, involving multiple injections of 6-OHDA administered at various sites within the pancreas. (B-C) Changes of body weight (B) and blood glucose (C) after sympathetic denervation. (D) Changes of blood glucose (left) and AUC (right) of DIO mice subjected to ipGTT. (E) Changes of serum insulin (left) and AUC (right) during ipGTT. (F) Changes of serum glucagon (left) and AUC (right) during ipGTT. (G)Changes of blood glucose (left) and AUC (right) of DIO mice subjected to ipITT. All data are mean ± SEM and analyzed by two-way ANOVA with multiple comparisons and unpaired t test. *p < 0.05, **p < 0.01.

We also performed cPSD in *db/db* mice (Figure. 6A). Compared to the vehicle group, neither body weight nor blood glucose was altered after 6-OHDA injection (Figure. 6B and 6C). Remarkably, *db/db* mice injected with 6-OHDA showed a higher glucose tolerance during GTT (Figure. 6D). Consistently, there was an increase in the accumulation of insulin secretion in response to glucose challenge (Figure. 6E). But there was a discernible downward trend observed in the accumulation of glucagon (Figure. 6F). Moreover, insulin sensitivity exhibited a significant enhancement after 21 days following 6-OHDA injection (Figure. 6G).

**Figure 6.**
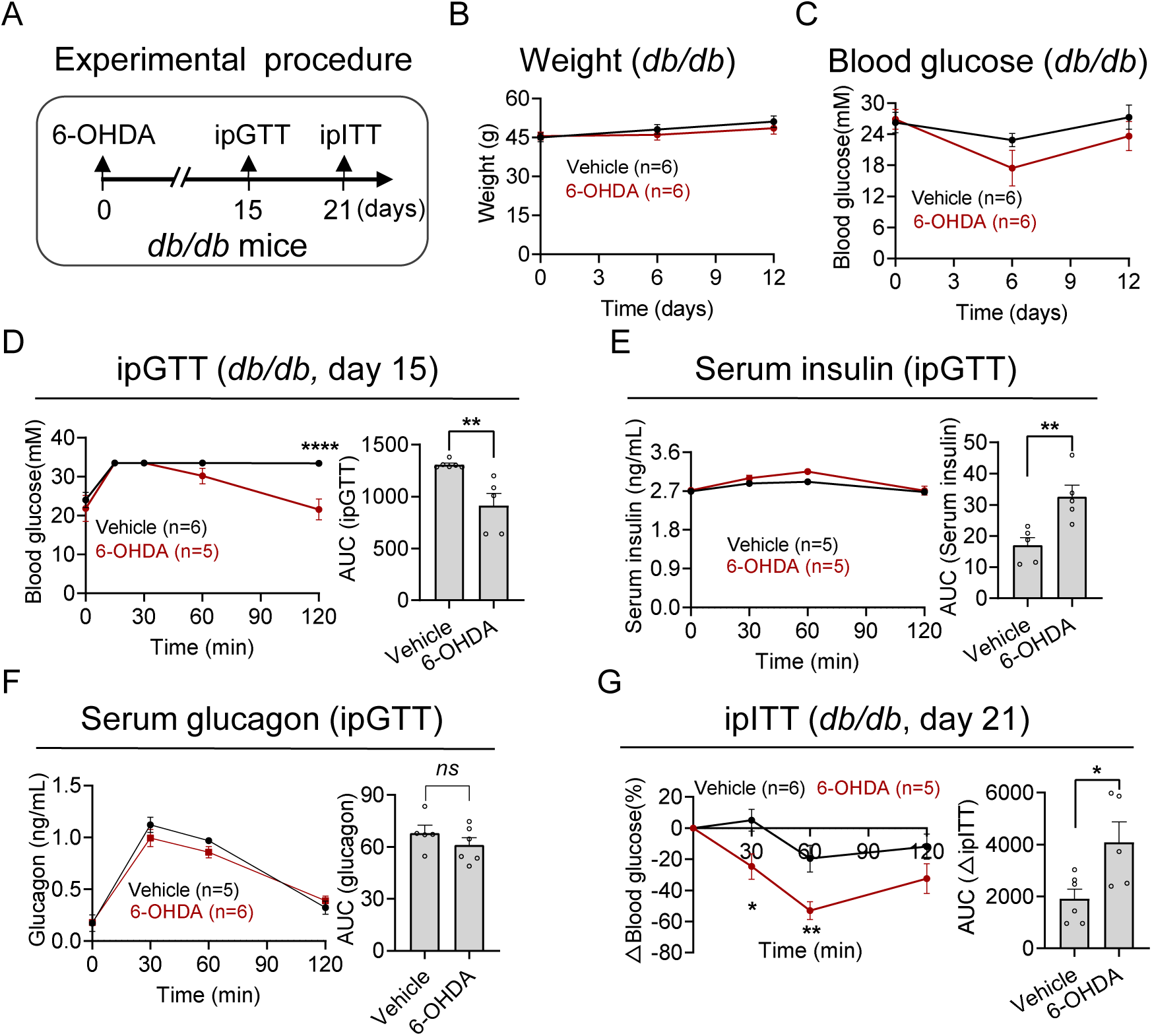
Effects of pancreatic sympathetic denervation in *db/db* mice. (A) An experimental design was implemented to examine the impact of sympathetic denervation on *db/db* mice (10-week-old), involving multiple injections of 6-OHDA administered at various sites within the pancreas. (B-C) Changes of body weight (B) and blood glucose (C) after sympathetic denervation in *db/db* mice. (D) Changes of blood glucose (left) and AUC (right) of *db/db* mice subjected to ipGTT. (E) Changes of serum insulin (left) and AUC (right) during ipGTT. (F) Changes of serum glucagon (left) and AUC (right) during ipGTT. (G) Changes of blood glucose (left) and AUC (right) of *db/db* mice subjected to ipITT. All data are mean ± SEM analyzed by two-way ANOVA with multiple comparisons and unpaired t test. *p < 0.05, **p < 0.01.

Taken together, cPSD results in an improvement in glucose tolerance in WT and *db/db* mice, but a decrement in DIO mice. Meanwhile, cPSD enhances insulin sensitivity in diabetic mice, with no observed impact on WT mice. These results suggested that pancreatic sympathetic innervation plays an important role in glucose metabolism, which is dependent on the physiological or pathological condition.

## Discussion

Sympathetic innervation and morphological changes in α/β-cells play pivotal roles in islet involvement in blood glucose regulation. Nevertheless, the intricate patterns of sympathetic innervation within islets remain poorly understood. To address this gap in knowledge, we meticulously analyzed the sympathetic innervation patterns in islets and their temporal dynamics utilizing high-resolution imaging and advanced 3D reconstruction techniques in adult WT mice, DIO mice, and *db/db* mice. Here, we offered a unique opportunity to identify anatomical and functional features of pancreatic sympathetic innervation in mice under different physio-pathological conditions, providing new strategy to improve diabetes. Accumulating evidence from laboratory and human studies suggested that therapeutically targeting sympathetic overactivity could help to prevent metabolic diseases [30]. Similar to the established use of renal sympathetic denervation in clinical practice for humans [31], we also look forward to the implement sympathetic intervention techniques for the management of diabetes and other pancreatic-related diseases.

### Alterations in the structural characteristics of pancreatic islets in response to diabetic condition

Employing advanced imaging methodologies, we have successfully delineated distinctive alterations in the distribution and composition of α/β-cells, sympathetic innervation within pancreatic islets of WT and diabetic mice (Figure. 1-3). In our study, we categorized islet size using systematic classification methods [29, 32]. By integrating data pertaining to islet size alongside its cellular constituents, we were able to provide a more nuanced depiction of the islet’s intrinsic microcellular milieu. We found small islets constituted the majority in pancreas of WT mice, and the percentage of small islets decreased significantly with aging (Figure. 2C). Besides, a tendency towards a decrease in islet sizes was noted in DIO mice (Figure. 2C), and a significant reduction was observed in *db/db* mice (Figure. 3B). Taken together, our findings suggest a propensity for pancreatic islets to undergo gradual enlargement with aging and elevated blood glucose levels, corroborating previous report [33]. This enlargement may originate from the expansion of β-cells mass, potentially initiated by downstream signaling of the insulin receptor or indirectly modulated through regulation of the peroxisome proliferator-activated receptor-gamma (PPAR-γ) [34, 35]. Concurrently, our findings indicated a positive correlation between aging or HFD intake and the enlargement of pancreatic islets. This suggests that an increase in the number of larger islets may be associated with a decline in cellular function, which is consistent with prior reports [36, 37]. Moreover, we noted that the ratio of α-: β-cells decreased with aging in both WT and DIO mice (Figure. 2E), but increased in *db/db* mice (Figure. 3D), indicating a desynchronization in the alterations of α-cells and β-cells as the islets undergo enlargement.

In WT mice, pancreatic islets exhibit a distinctive arrangement where β-cells predominantly located in the core of islets, surrounded by α-cells in the shell (Figure. 1A), consistent with prior reports [7, 38, 39]. With advancing age in mice, there is an increase in the number of α-cells within the core region (Figure. 2D). Interestingly, a distinct distribution pattern of α-cells and β-cells is evident in the islets of *db/db* mice (Figure. 3C). Unlike the core-mantle structure observed in WT and DIO mice (Figure. 1A and 2D), α-cells and β-cells are intermixed in *db/db* mice (Figure. 3C). This unique distribution, similar as in human [11, 39, 40], bears implications for the distinct functional role of α/β-cells in the diabetic condition. Notably, there was an increase in the distribution of α-cells within the core of islets in *db/db* mice, suggesting a potential compensatory mechanism in response to the diabetic milieu. Our findings enhance the comprehension of the intricate organization and functional implications of pancreatic islets in different genetic backgrounds and dietary conditions. This comprehensive analysis contributes to the improvement of clinical protocols for islet transplantation, thereby propelling progress across a spectrum of endeavors in the fields of islet and pancreatic research and therapy.

### Differences of pancreatic sympathetic innervation between WT and diabetic mice

Compared to WT-10 mice, the proportion of TH-innervated β-cells over total β-cells increased in WT-26 mice (Figure. 2G), indicating that aging is a factor influencing the sympathetic innervation in islets. In DIO mice, both TH-innervated α-cells and β-cells exhibited reductions (Figure. 2F and 2G), suggesting HFD affect the pancreatic sympathetic nerves. However, although TH-innervated α-cells decreased (Figure. 3E), the TH-innervated β-cells increased in *db/db* mice (Figure. 3F). A previous study reported an inclination towards increased noradrenergic innervation of the endocrine area in obese *db/db* mice with aging as diabetes progresses [23]. This discrepancy could be attributed to factors such as excessively thin section thickness, outdated imaging modalities, and suboptimal software quantification. Our data revealed that islet α/β-cells possessed relatively abundant sympathetic innervation in both WT and diabetic mice. Furthermore, our study profoundly reinforces the notion that the sympathetic innervation of mouse islet β-cells is conspicuously elevated in comparison to prior reports [21, 22], thereby carrying momentous implications for sampling accuracy and cellular functionality.

In our histomorphological study, we observed TH-expressing cells within islets, with a reduction in diabetic mice. It is noteworthy that some of the TH^+^ cells exhibited distinctive dendritic structures (Figure. 1H), establishing contacts with endocrine cells. As reported [41–44], some β-cells expressed TH, which is crucial for insulin secretion in mice. Hence, these factors should be taken into consideration in future studies exploring these innervations and their function.

### Pancreatic sympathetic innervation in the glucose regulation under diabetic state

Philip Borden found no defects were observed in glucose tolerance of sympathectomized mice [7], but Jimenez-Gonzalez reported that activation of sympathetic neurons impaired glucose tolerance [45]. We found that glucose tolerance was greatly improved in WT mice with pancreatic sympathetic denervation via 6-OHDA in situ injection (Figure. 4E). Notably, both the glucose tolerance and insulin sensitivity has been significantly improved in *db/db* mice (Figure. 6D and 6G). Also, there was a marked increase in insulin sensitivity of DIO mice (Figure. 5G). The only discrepancy is that sympathetic denervation decreases glucose tolerance in DIO mice (Figure. 5D). The opposite effect on ipGTT in the DIO and *db/db* mice might be attributed to genetic background since *db/db* mice are leptin receptor deficient.

Overall, our findings provide insights into the impact of sympathetic innervation on blood glucose regulation under different physio-pathological conditions. At present, most anti-diabetic drugs regulate blood sugar via enhancing the secretion of insulin or improving insulin sensitivity. As our results suggested, pancreatic sympathetic denervation improved the insulin sensitivity in both DIO and *db/db* mice. We believe that the pancreatic sympathetic modulation might become a novel therapeutic strategy for T2D, resembling the renal denervation for the patients with hypertension [31].

### Limitations of the study

In this study, we explored the roles of pancreatic sympathetic nerves in physiological and diabetic conditions, suggesting that the potential of manipulating pancreatic sympathetic nerves as a novel therapeutic approach for T2D. However, given both α-cells and β-cells in the pancreas are innervated by sympathetic nerves, it was challenging to selectively manipulate nerves that only target α-cells or β-cells. Further studies should be directed towards in-depth analyses of sympathetic nerves, focusing on uncovering their molecular profiles, cellular properties, and the dynamics roles they play.

## Note

Our article has been submitted by *Acta Biochimica et Biophysica Sinica* (ABBS).

## Statement

Animal experiment was approved by the Laboratory Animal Management Committee of ShanghaiTech University (approval Number. 20230809001).

## Acknowledgments

We thank Dr. Xiaoming Li and the Molecular Imaging Core Facility (MICF) of School of Life Science and Technology, ShanghaiTech University for Microscopic imaging; and the staff members of the Animal Facility at ShanghaiTech University for providing technical support and assistance.

---

**Table S1.** Key resources table.

| REAGENT or RESOURCE | SOURCE | IDENTIFIER |
| --- | --- | --- |
| **Antibodies** |  |  |
| Chicken anti-TH | Sigma/Aldrich | AB9702 |
| Mouse anti-Insulin | Cell Signaling | L6B10 |
| Rabbit anti-Glucagon | Cell Signaling | 2760 |
| Goat anti-rabbit IgG, Alexa Fluor 488-conjugated | Jackson | 111-545-003 |
| Goat anti-Mouse IgG, Alexa Fluor 568-conjugated | Invitrogen | A11031 |
| Goat anti-chicken IgG, Alexa Fluor 647-conjugated | Invitrogen | A-21449 |
| **Chemicals, peptides, and recombinant proteins** |  |  |
| DAPI | Beyotime | P0131 |
| stypic powder | KELC | N/A |
| QuickBlock™ Blocking Buffer | Beyotime | P0260 |
| QuickBlock™ Primary Antibody Dilution Buffer | Beyotime | P0262 |
| QuickBlock™ Secondary Antibody Dilution Buffer | Beyotime | P0265 |
| **Software and algorithms** |  |  |
| GraphPad Prism 8 | GraphPad Software | https://www.graphpad.com/scientific-software/prism/ |
| Imaris 9.7.2 | Imaris | N/A |
| Fiji, ImageJ | NIH | https://imagej.net/Fiji |
| Huygens Professional 19.4 | Huygens | N/A |
| ZEN2012 black/blue edition | Zeiss | N/A |
| Leica Application Suite X (TCS SP8) | Leica | Wetzlar, Germany |
| Nikon CSU-W1 Sora 2 Camera | Nikon | Japan |
| OlyVIA Vs120 | Olympus | N/A |

**Figure S1.**
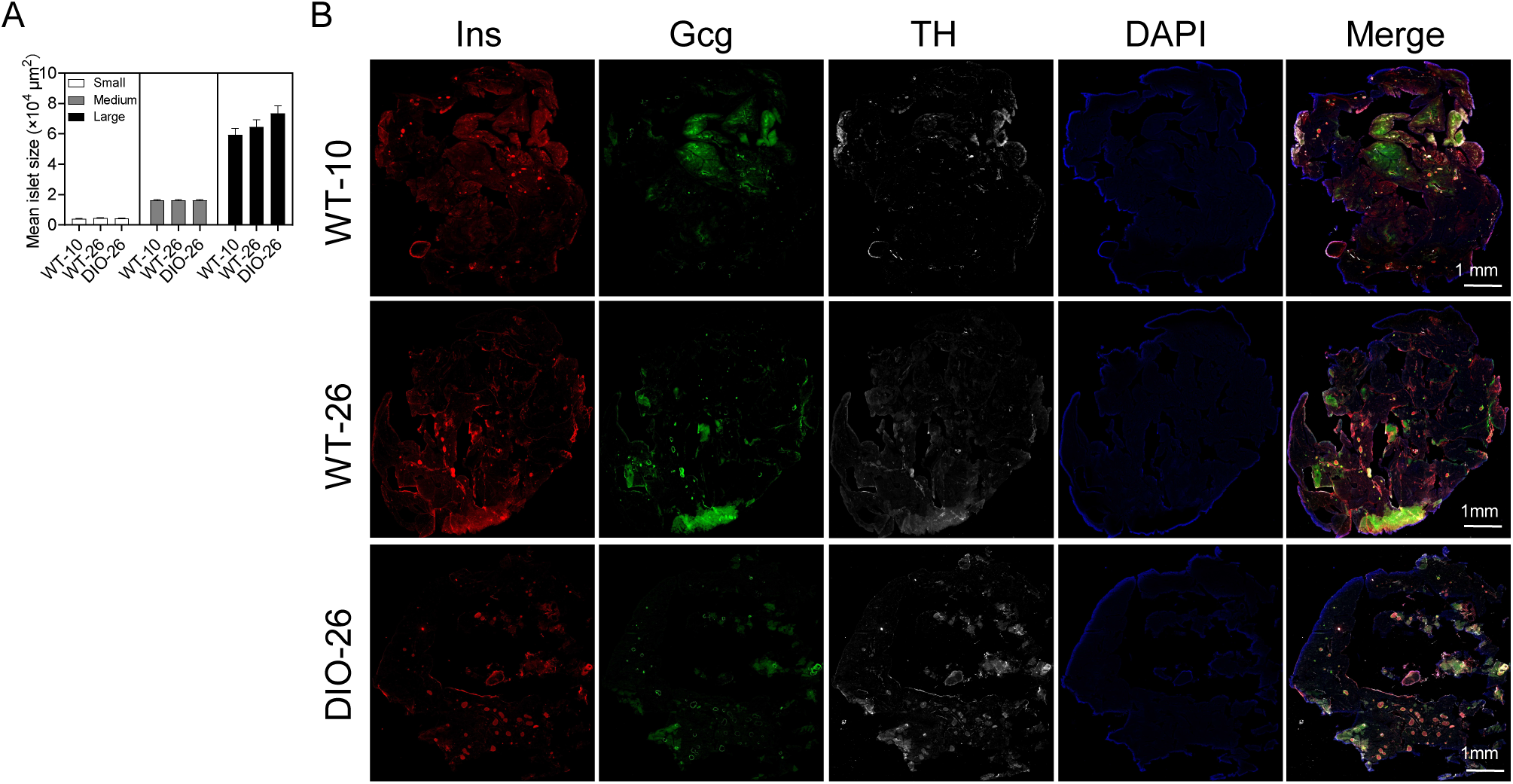
Islets in DIO mice, related to Fig. 2. (A) Average area of islets with different sizes in WT and DIO mice. (WT-10/26 refers to WT mice of 10 weeks old and 26 weeks old respectively, DIO-26 refers to WT-10 mice fed with HFD for 16 weeks) (B) Overview of pancreatic slice sections in WT and DIO mice.

**Figure S2.**
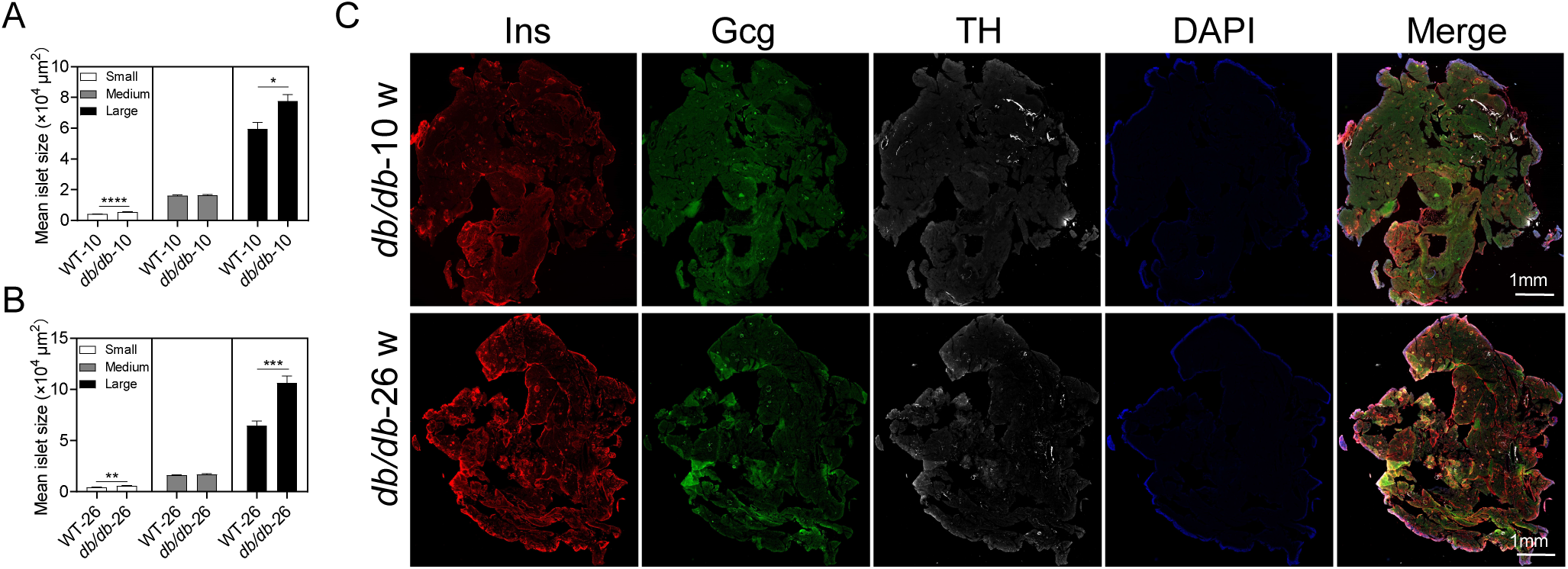
Islets in db/db mice, related to Fig. 3. (A-B) Average area of islets with different sizes in WT mice and db/db mice aged 10 weeks old (A) and 26 weeks old (B). (C) overview of pancreatic slice sections in db/db mice.

